# Capturing Optimal and Suboptimal behavior Of Agents Via Structure Learning Of Their Internal Model

**DOI:** 10.1101/2024.09.30.615767

**Authors:** Ashwin James, Ingrid Bethus, Alexandre Muzy

**Affiliations:** Université Côte d’Azur - CNRS - I3S; Université Côte d’Azur - CNRS - IPMC

## Abstract

This study introduces a novel framework for understanding the cognitive underpinnings of individual behavior which often deviates from rational decision-making aimed at maximizing rewards in real-life scenarios. We propose a structure learning approach to infer an agent’s internal model, composed of a learning rule and an internal representation of the environment. Crucially, the combined contribution of these components — rather than their individual contribution — determines the overall optimality of the agent’s behavior. By exploring various combinations of learning rules and environment representations, we identify the most probable agent model structure for each individual. We apply this framework to analyze rats’ learning behavior in a free-choice task within a T-maze with return arms, evaluating different internal models based on optimal and suboptimal learning rules, along with multiple possible representations of the T-maze decision graph. Identifying the most likely agent model structure based on the rats’ behavioral data reveals that slower learning rats employed either a suboptimal or a moderately optimal agent model, whereas fast learners employ an optimal agent model. Using the inferred agent models for each rat, we explore the qualitative differences in their individual learning processes. Policy entropy, derived from the inferred agent models, also highlights variations in the balance between exploration and exploitation strategies among the rats. Traditional reinforcement learning approaches to addressing suboptimal behavior focus separately on either suboptimal learning rules or flawed environment representations. Our approach jointly models these components, revealing that suboptimal decisions can arise from complex interactions between learning rules and environment representations within an agent’s internal model. This provides deeper insights into the cognitive mechanisms underlying real-world decision-making.

## 1 Introduction

In real-world scenarios, individuals often exhibit irrational or suboptimal behavior, making choices that do not maximize their rewards, unlike an ideal rational agent (Kahneman and Tversky, 2013; Sharot et al., 2007; O’Donoghue and Rabin, 2000; Zentall, 2015). This results in decisions or actions that fail to achieve the best possible outcome given the available information and constraints (Kahneman, 2003; Rubinstein, 1998). Studies have shown that deviation from rational behaviour may occur due to cognitive biases (Evans, 1989; Zentall, 2019), heuristic decision strategies (Evans, 1984; Gigerenzer and Gaissmaier, 2011), emotional stress (Starcke and Brand, 2012) or mental illness (Mukherjee and Kable, 2014) among many other causes. In reinforcement learning (RL) models of individual behavior, suboptimality is typically handled in two ways: as irrational learning rule or a flawed internal representation of the environment. A suboptimal learning rule hinders an agent’s ability to effectively learn from experiences, leading to choices that may not maximize rewards. Conversely, an inefficient or flawed internal environment model can impede the learning process by providing an inaccurate or incomplete representation of the world (Solway et al., 2014). It’s also important to note that the interplay between learning rules and environment models is complex. While a particular learning rule might be effective with one environment model, it could be inefficient with another.

To capture the underlying dynamics of individual behavior, we propose a structure learning framework that can learn the internal model of an individual agent. Specifically, by “structure”, we refer to the components of an RL agent’s internal model, which include a learning rule (selected from a range of suboptimal and optimal rules) and an environment representation (chosen from various representations). Our framework infers the agent’s internal model, which includes both a learning rule and an internal environment representation, from a pool of possible agent models consisting of various learning rules and environment representations.

We apply our framework to a free-choice task in which rats navigate a T-maze with returning arms, receiving rewards for following predefined correct paths to two feeder boxes. We infer the agent’s structure by identifying the combination of learning rule and environmental representation that maximizes the Bayesian Information Criterion (BIC) score from a predefined set of agent models. We define rats’ strategies as the internal agent models employed by them, which are given by the combination of their learning rules and internal environment representation.

While Reinforcement Learning (RL) typically assumes that agents behave optimally, aiming to maximize their returns, human and animal behavior often deviates from this ideal behaviour due to cognitive biases (e.g., the sunk cost fallacy, where past investments influence future choices (Magalhães et al., 2012; Sweis et al., 2018)). Simple heuristics, even if seemingly “irrational”, can sometimes better explain individual behavior (Hayden and Niv, 2021). For instance, individuals might stick to past successful choices without exploring new options, relying on heuristics. To analyze rats’ learning behaviour in the maze experiment, we employ two learning rules: an optimal learning rule represented by the Temporal Difference (TD) based Q-learning with an eligibility trace, which we refer to as Discounted RL (DRL) algorithm (Sutton and Barto, 2018) and a suboptimal heuristic-based learning rule called Cognitive Activity-based Credit Assignment (CoACA) (James et al., 2023; Muzy, 2019).

TD learning has been extensively used to address the credit assignment problem in animal learning (Niv, 2009; Maia, 2009; O’Doherty et al., 2015; Samson et al., 2010; Dolan and Dayan, 2013; Schultz et al., 1997; Bathellier et al., 2013). The DRL algorithm aims to maximize expected future returns, aligning with the concept of economically rational behavior and characterizing optimal agents. However, since animal and human behavior is not always optimal, we propose CoACA as a simpler heuristic alternative to DRL that prioritizes action duration within rewarding sequences, potentially leading to suboptimal behavior. Agents using CoACA assign larger credits to actions with longer durations because these actions are more memorable (James et al., 2023). This approach is considered suboptimal compared to DRL-based agents, which aim to maximize expected returns and are therefore deemed optimal. Figure 1a presents a Semi-Markov Decision Process (SMDP) model of an RL problem where agents have an internal model composed of a learning rule and an internal environment representation, with each action having a random action duration.

**Figure 1:**
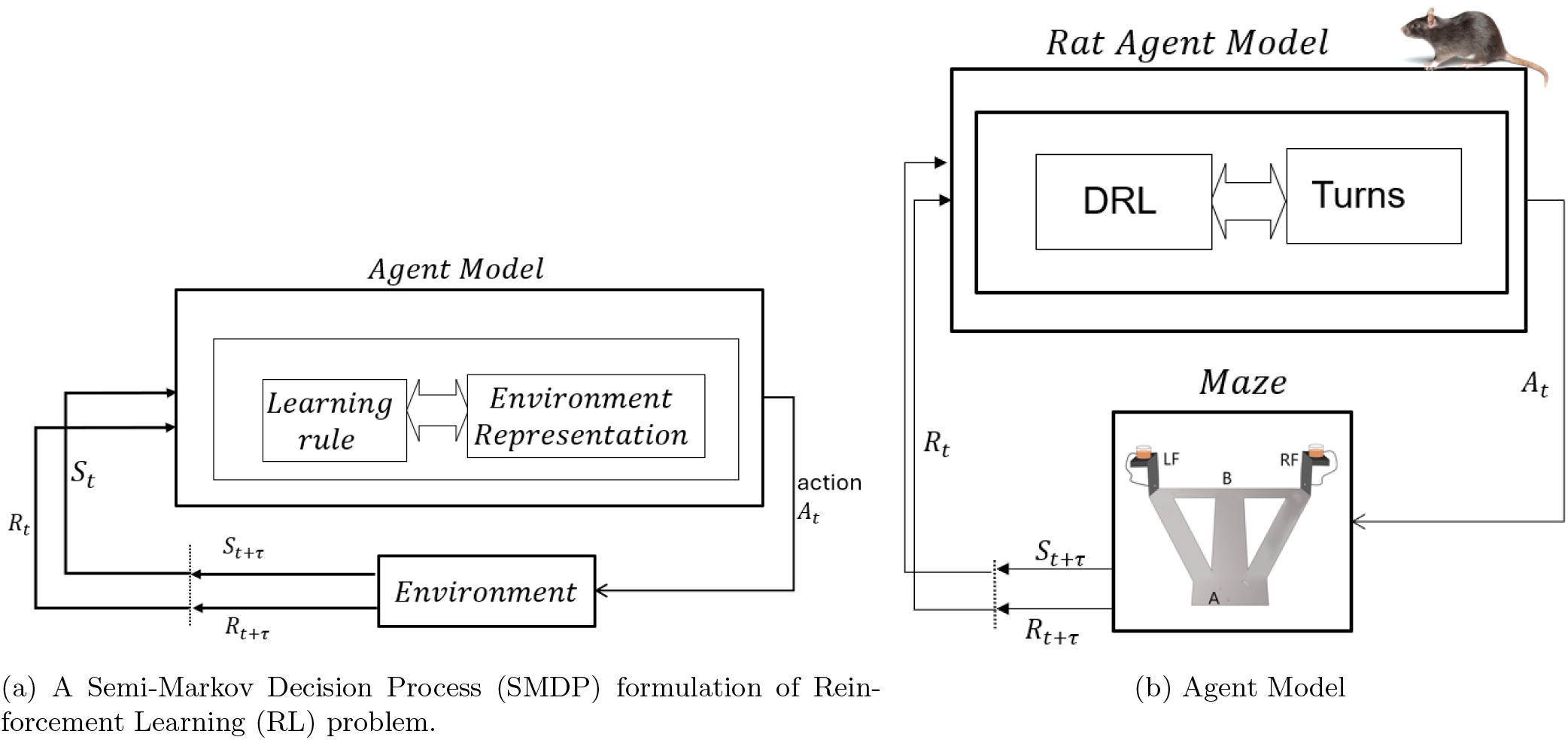
(a) Semi-Markov Decision Process (SMDP) (see Section 4.1) with Agent Model: an SMDP representation of RL problem where an agent, whose internal model is given by a combination of learning rule and an internal environment representation, chooses actions that have random durations given by *τ* . (b) A possible agent model structure for rat experiment, where the learning rule is given by DRL and the environment representation is a Turns model of the maze.

To define the environment models in the agents’ internal representations, we created six decision graphs (see Figure 3) that represent the rats’ internal maps of the maze used in the experiment (see Figure 2a). These models vary from highly detailed to more abstract representations of the environment. The Turns model provides the most granular view by breaking the maze into its smallest segments between each decision point, whereas the Paths model offers a simplified overview by combining all atomic segments between the feeder boxes into a single path. The four “hybrid models”, positioned between these extremes, integrate elements of both the Turns and Paths models, incorporating different levels of details. (Figure 1b represents a possible agent model of rats).

**Figure 2:**
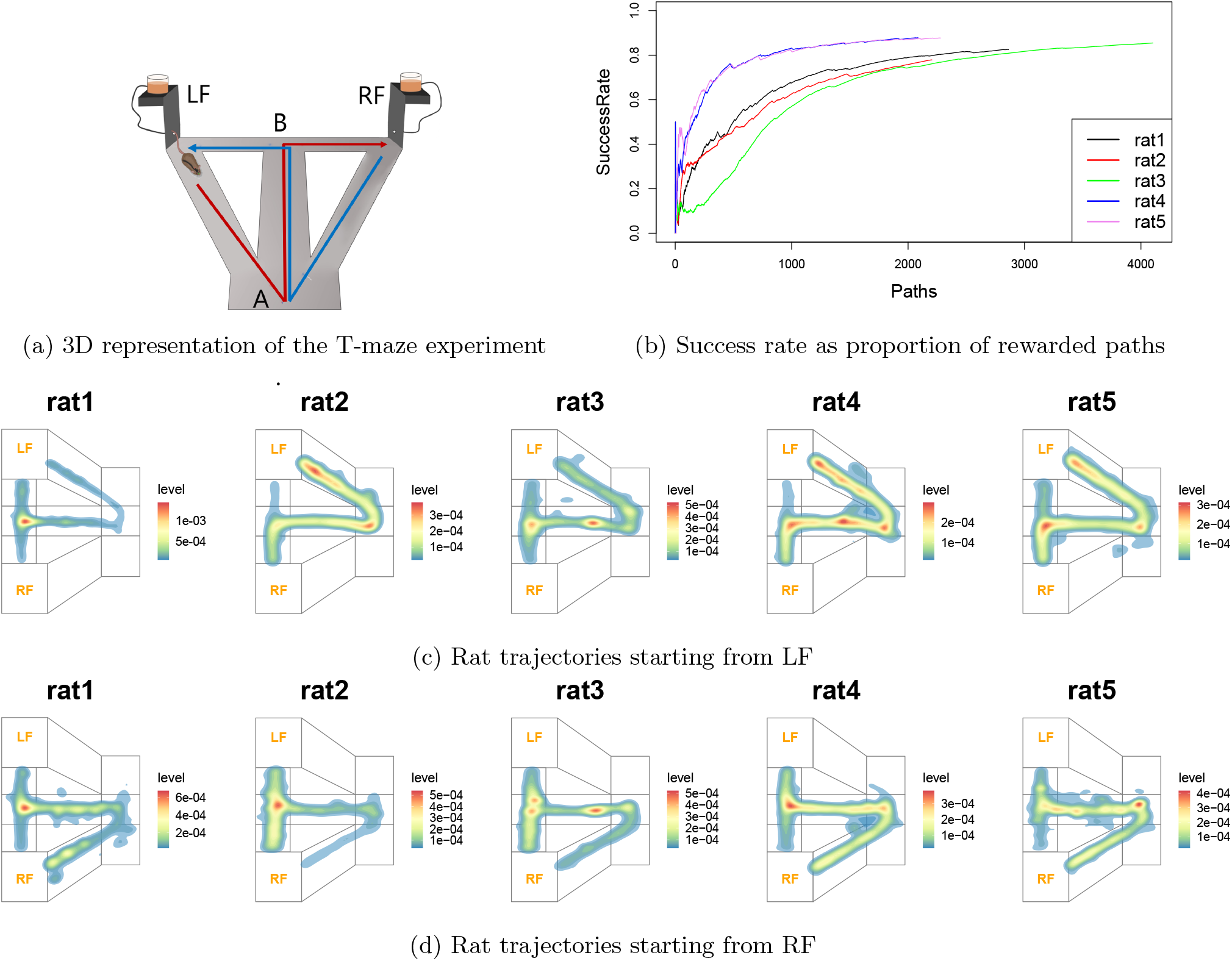
Maze and raw data analysis: (a) 3D representation of the T-maze exp: A, B, LF and RF are four choice points; LF and RF represent the left and right feeders respectively. Good path from LF, Good.LF, is shown in red, while good path from LF, good.RF, is shown in blue. (b) Success rate computed as proportion of rewarded paths: The rats can be categorized as slower learning (rat1, rat2, rat3) or faster learning (rat4, rat5) based on the proportion of rewarded paths. (c,d) Heatmap comparing the trajectories of rats during the learning phase (first 400 paths). Levels represent the density of rat positions observed at each position in the maze, computed by two-dimensional kernel density estimation with an axis-aligned bivariate normal kernel (Venables and Ripley, 2002) using R library Modern Applied Statistics with S (MASS)^1^. **rat1**: *LF trajectories do not indicate a pattern, RF shows high density over Good*.*RF path*. **rat2**: *LF trajectories have high density for Good*.*LF path, while RF shows high density for Straight*.*RF*. **rat3**: *Same as rat2*. **rat4**: *High density for Good*.*LF and Good*.*RF paths*. **rat5**: *Same as rat4*. Slow learning rats (rat1, rat2, rat3) learn the good path from one feeder while mostly preferring the straight path from the other feeder. Conversely, the faster learning rats’ heatmaps show that they learn both good paths early during the experiment.

**Figure 3:**
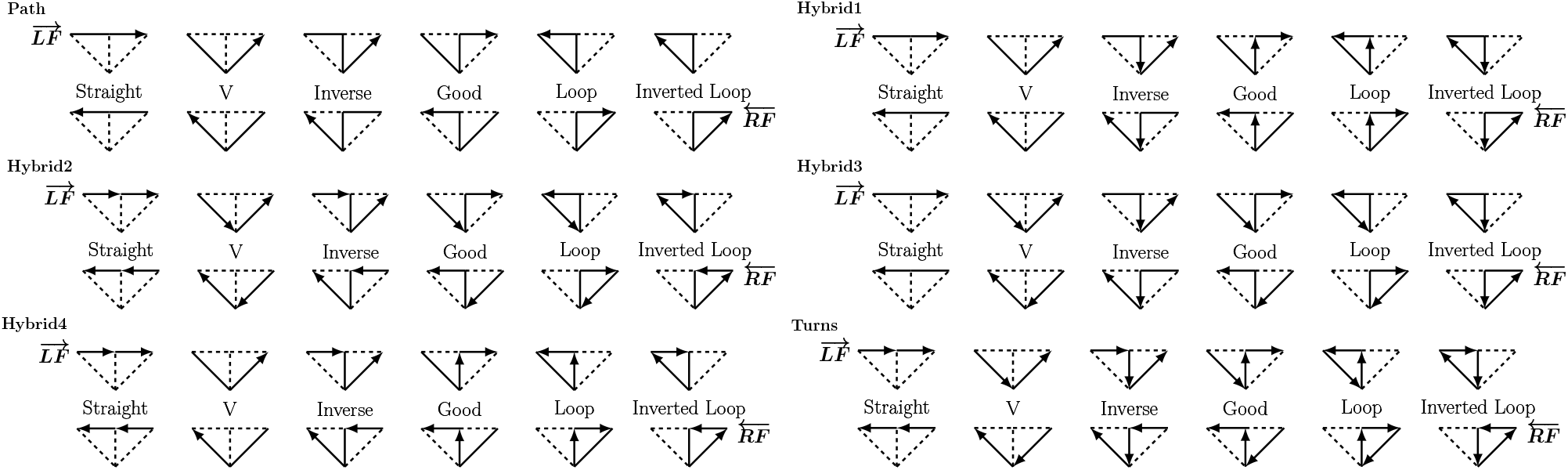
The six different cognitive graphs of the maze. Actions are depicted as arrows between environmental states: choice points A and B, Left Feeder (LF) and Right Feeder (RF) of the maze, as shown in Figure 2a.

While environment representations illustrate the various ways to internally construct an environment model, we hypothesize that the efficiency of the agent structure depends on the combination of the learning rule and environment representation. The Turns model, with its fine-grained structure, supports frequent TD updates and leads to faster learning compared to other models. Thus, for the maze task, the agent model combining DRL with Turns may be the optimal performer. Conversely, agent models based on CoACA are deemed suboptimal due to a forgetfulness factor that causes credit decay at the end of each session (as detailed in Section 4.1). The remaining agent models are moderately optimal, with performance variations largely driven by the intervals between decision points within each model.

Previous research has applied reinforcement learning (RL) models to study individual behavior in reward-based tasks (Morris et al., 2006; Roesch et al., 2007; Maia, 2009; Constantino and Daw, 2015). However, a major limitation of traditional RL models is their assumption that agents are rational and driven solely by reward maximization. Since human and animal behavior often deviates from this ideal, influenced by cognitive biases and suboptimal decision-making, various modifications to RL learning rules have been proposed to capture these suboptimal behaviors. These modifications typically introduce additional parameters or mechanisms to address factors such as pessimistic agents modeling anxiety disorders (Zorowitz et al., 2020), the “stickness” tendency to repeat previously chosen options regardless of their outcomes (Kanen et al., 2019; Suzuki et al., 2023), and differential processing of rewards and punishments (Lim et al., 2019). (Reddy et al., 2018) suggest that suboptimal behavior may stem from a flawed internal environment representation of agents, while (Kwon et al., 2020) propose that individuals might hold incorrect assumptions about the world, as reflected in their internal environment models.The approaches mentioned above independently capture individual suboptimality either through suboptimal learning rules or flawed internal environment models. In contrast, our framework allows jointly inferring both learning rule and internal environment representations as part of the agent’s internal model, where the optimality of agent models relies on the combination of these components, rather than on each component individually.

Applying the agent model structure learning framework to experimental data of rats identified two distinct learning profiles in rats. Slower learners were best characterized by a hybrid environment model combining either CoACA and DRL learning rules. In contrast, faster learners were optimally represented by an agent model combining Turns environment with DRL learning rule. The inferred agent structures were further used for analyzing qualitative differences among individuals in their learning behavior by measuring their policy entropies. Policy entropies showed that rats displayed varying levels of explorative (high policy entropy) and exploitative (low policy entropy) behaviors during the learning stage. Additionally, individual responses to policy entropy differed: some rats took longer time to choose actions when faced with high entropy (higher uncertainty), while others reduced their choice duration. The variability in the inferred agent models, coupled with these finer qualitative differences in rats’ behaviors, underscores the complexity of individual learning behavior in natural intelligence.

While traditional RL methods focus on learning identifying “high-value” choices that maximize rewards, animals and humans exhibits greater sophistication, including the ability to construct causal world models (Lake et al., 2017). These models capture the causal relationships between actions, states, and outcomes, explaining how states evolve with actions and predicting the expected rewards from those transitions. However, current RL approaches often overlook the critical role of an agent’s internal environment representation (causal world model) in shaping individual behavior (Ashwood et al., 2020; Kwon et al., 2020; Ashwood et al., 2022; Schultheis et al., 2021). Our work addresses this limitation by introducing a novel framework that integrates RL methods with causal world models. By combining a learning rule with an environment representation in an internal agent model, we offer a more comprehensive approach to capturing the interplay between an individual’s learning process and their understanding of the world, thereby deepening our understanding of the cognitive foundations of real-life decision-making.

## 2 Results

### 2.1 Experiment and behavioral raw data analysis

Five male Long-Evans rats were used in the experiment. To motivate the rats to collect food rewards from the maze, they were subjected to a food deprivation program by keeping them at 90% of their body weight during the experiment. Each rat has multiple sessions in the maze, where each session lasts 20 minutes. During the sessions, the rats can freely move around in the maze uninterrupted. The T-maze with return arms (*cf*. Figure 2a) has two feeder places, Left Feeder (LF) and Right Feeder (RF), where the rats could get a food reward. The maze consists of a central stem (100*cm* long), two choice arms (of 50*cm* each) at one end of the central stem and two lateral arms connecting the other end of the central stem to the choice arms.

The 5 rats can be roughly classified into two groups based on their learning speed: rat1, rat2 and rat3 take more time to learn the two good paths compared to rat4 and rat5 that are faster learners. As depicted in Figure 2b, this difference can be seen in the plot of the proportion of rewarded paths made by the rats.

The heat maps of the rats’ trajectories during the first 400 paths show that the faster-learning rats have mastered both reward paths in the maze, while the slower-learning rats have not yet achieved this. During this learning stage, faster learning rats, 4 and 5, have a higher density over the good path trajectories from both feeders(*cf*. Figure 2). In case of rat1, Figure 2d shows that it has possibly learned the good path from RF, while LF heatmap does not show any particular pattern. For rat2 and rat3, the LF heatmap, shown in Figure 2d, has high density over good path trajectory indicating that they have learned the good path from LF, while the RF heatmap shows high density over the straight path from RF to LF.

### 2.2 Possible agent structures to model rats’ behaviors

Rats being free to move, there is no clear beginning and end of the task for the animal. They have to infer over time their own credit assignment to actions based on their own maze representation. Over credit assignment, rats can choose between different temporal representations of the reward consequences of their actions on the environment.

For each credit assignment method, either DRL or CoACA, the possible maze representations of the rat can be constructed in different ways by grouping the actions at choice points A and B. The assumption in these graphs is that the rats are always aware of three variables: “last visited feeder box”, “last chosen action” and “current head position”. The trajectories of the rats can be described by six different actions starting from each feeder where the good path is the only one leading to a reward (*cf*. Figure 3). Figure consists of six different cognitive graphs of the maze with different degrees of choice granularity. The Path model consists of single actions that takes the rat from one feeder to another/back to the same. Turn model represents the other extreme of choice granularity where the rat is required to always choose actions at choice points A and B. The Hybrid models describe chunked versions of actions from the Turn model: (i) Hybrid1: rat selects an action at the second choice, not the first, (ii) Hybrid2: rat selects an action at the first choice point, but not the second one, (iii) Hybrid3: rat selects an action only at choice point A, (iv) Hybrid4: rat selects an action at choice point B.

In DRL, the credit (value) of an action, as the result of choosing an action, is discounted proportionally to the delay to the future reward. DRL thus discounts future rewards and thereby gives more weight to immediate rewards. DRL credit assignment leads implicitly to *action duration reduction*, as it can be seen in Equation 7 .

Unlike DRL, CoACA does not consider future payoffs, but instead reinforces persistence of action choices by accumulating their past credits (*cf*. Section 4.1). This credit depends on both behavioral activity and performance of animals. This allows CoACA to propose a theory of activity-based credit assignment that better assesses the dynamics of different action choice strategies during the learning, while also offering a plausible explanation for suboptimal behavior. To do so, at the end of each episode, the activity-based credit of an action is computed based on its behavioral activity (duration relative to the duration of the episode) multiplied by the number of rewards accumulated in the episode, 0, 1 or 2. Note that while DRL credit assignment lead implicitly to *action duration reduction*, CoACA accounts for *action duration contribution*, here, the different action durations in the different maze representations of the rats.

### 2.3 Agent Model Recovery From Simulated Data

To test the recovery of agent models, we generated simulated data with 50 simulations for each of the 12 different agent models (AMs) using model parameters identified from rats’ experimental data and attempted to recover the true generated AM. The confusion matrix representing *Pr*(fit model | simulated model) is shown in Table 1 and demonstrates that inferring the true model based on minimum BIC scores achieves a good recovery rate for all AMs.

**Table 1:** Confusion Matrix showing the agent model recovery rates. AM1 = Paths.CoACA, AM2 = Hybrid1.CoACA, AM3 = Hybrid2.CoACA, AM4 = Hybrid3.CoACA, AM5 = Hybrid4.CoACA, AM6 = Turns.CoACA, AM7 = Paths.DRL, AM8 = Hybrid1.DRL, AM9 = Hybrid2.DRL, AM10 = Hybrid3.DRL, AM11 = Hybrid4.DRL, AM12 = Turns.DRL

|  | AM1 | AM2 | AM3 | AM4 | AM5 | AM6 | AM7 | AM8 | AM9 | AM10 | AM11 | AM12 |
| --- | --- | --- | --- | --- | --- | --- | --- | --- | --- | --- | --- | --- |
| AM1 | 1.0 | 0.0 | 0.0 | 0.0 | 0.0 | 0.0 | 0.0 | 0.0 | 0.0 | 0.0 | 0.0 | 0.0 |
| AM2 | 0.0 | 1.0 | 0.0 | 0.0 | 0.0 | 0.0 | 0.0 | 0.0 | 0.0 | 0.0 | 0.0 | 0.0 |
| AM3 | 0.0 | 0.0 | 1.0 | 0.0 | 0.0 | 0.0 | 0.0 | 0.0 | 0.0 | 0.0 | 0.0 | 0.0 |
| AM4 | 0.0 | 0.0 | 0.0 | 1.0 | 0.0 | 0.0 | 0.0 | 0.0 | 0.0 | 0.0 | 0.0 | 0.0 |
| AM5 | 0.0 | 0.0 | 0.0 | 0.0 | 1.0 | 0.0 | 0.0 | 0.0 | 0.0 | 0.0 | 0.0 | 0.0 |
| AM6 | 0.0 | 0.0 | 0.0 | 0.0 | 0.0 | 1.0 | 0.0 | 0.0 | 0.0 | 0.0 | 0.0 | 0.0 |
| AM7 | 0.0 | 0.0 | 0.0 | 0.0 | 0.0 | 0.0 | 1.0 | 0.0 | 0.0 | 0.0 | 0.0 | 0.0 |
| AM8 | 0.0 | 0.0 | 0.0 | 0.0 | 0.0 | 0.0 | 0.0 | 0.98 | 0.0 | 0.0 | 0.0 | 0.02 |
| AM9 | 0.0 | 0.0 | 0.0 | 0.0 | 0.0 | 0.0 | 0.0 | 0.0 | 1.0 | 0.0 | 0.0 | 0.0 |
| AM10 | 0.0 | 0.0 | 0.0 | 0.0 | 0.0 | 0.0 | 0.0 | 0.0 | 0.02 | 0.98 | 0.0 | 0.0 |
| AM11 | 0.0 | 0.0 | 0.0 | 0.0 | 0.0 | 0.0 | 0.0 | 0.0 | 0.02 | 0.0 | 0.98 | 0.0 |
| AM12 | 0.0 | 0.0 | 0.0 | 0.0 | 0.0 | 0.0 | 0.0 | 0.08 | 0.0 | 0.02 | 0.0 | 0.90 |

### 2.4 Inferring the best fitting agent structure

To select the most appropriate credit assignment for each rat, 12 different models were compared, constructed by applying the credit assignments CoACA and DRL to each of the six maze representations: Paths, Hybrid1, Hybrid2, Hybrid3, Hybrid4 and Turns (*cf*. Figure 3). The results of likelihood comparison are shown in Table 2. More details about the selection method are provided in Section 4.2.

**Table 2:**
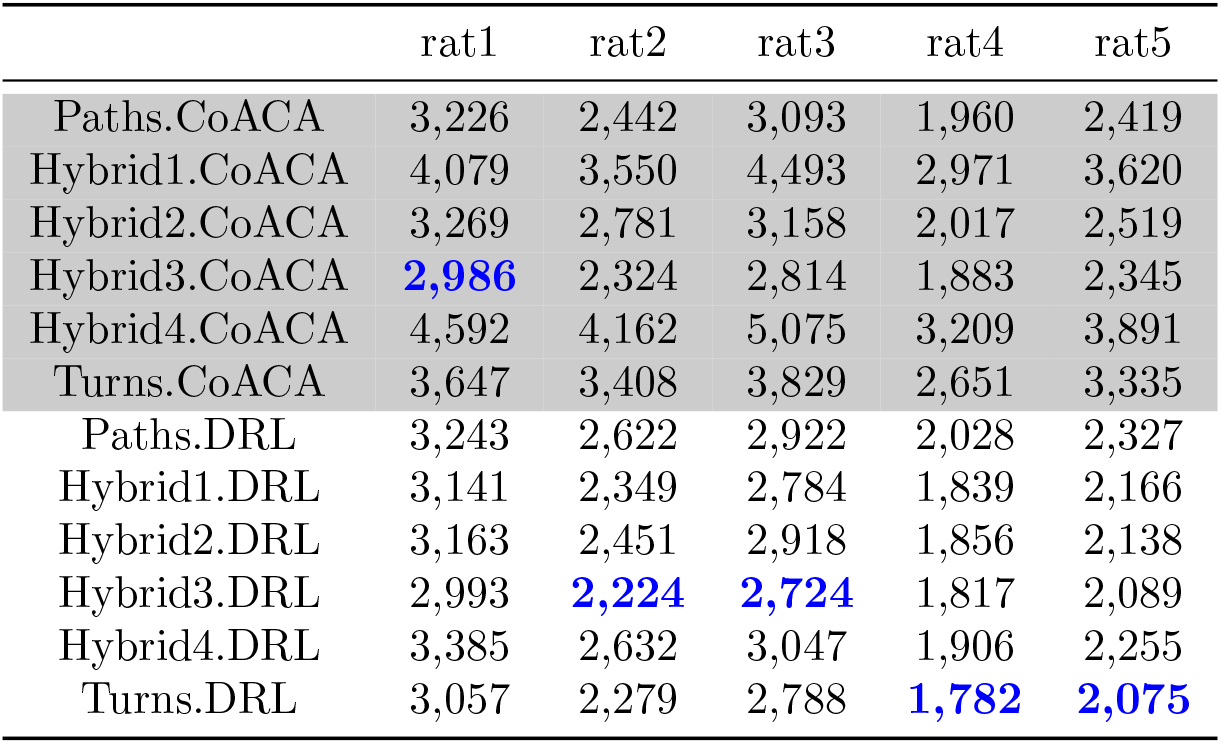
Bayesian Information Criterion (BIC) scores computed for each rat and each model.

Individual rat behaviors varied significantly as reflected by their best-fitting agent model structures. Rat 1 exhibited slower learning and was best modeled by CoACA.Hybrid3, while rats 2 and 3, also slow learners, were better represented by DRL.Hybrid3. In contrast, rats 4 and 5, characterized by faster learning, were best modeled by DRL.Turns, where the Turns model of the maze with its higher frequency TDCA updates, captured better the accelerated learning in the fast-learning rats.

### 2.5 Policy Entropy Of Individuals Based On Their Inferred Agent Models

The most likely agent model structures from Table 2 for each rat are used to construct the rats’ policies through the course of the experiment. Entropy of the policies at each path is computed using Equation 13. Entropy measure can be high or low - high policy entropy indicates that the rats have not yet learned which action to select in a given state. This implies that the rats are still exploring for the optimal behavior, resulting in less structured behavior where any action could occur in any state. In contrast, low entropy signifies low uncertainty. This means that the rats have learned which action to choose in a given state, leading to a clear and consistent pattern of actions within each state. The entropy plots show roughly two different behaviors:

1. **Low Entropy Policies:** Rat2, rat4, and rat5 exhibit more consistent policies based on Hybrid3.DRL, with steadily declining entropy until the rats learn the optimal behavior. This indicates more structured behavior with less variability, akin to exploitative behavior.
  - For rat2, Hybrid3.DRL predicts a low entropy policy during the learning stage, even though rat2 follows a suboptimal policy by learning the good path in LF and taking the straight path from RF (cf. Figure 4).
  - For rats 4 and 5, Turns.DRL predicts low-entropy policies (cf. Figure 4).
2. **High Entropy Policies:** The policies of rat1 (based on Hybrid3.CoACA) and rat3 (based on Hybrid3.DRL) display a high level of entropy during the learning stages (approximately the first 600 paths). Additionally, Hybrid3.CoACA predicts not only high policy entropy in action selection but also greater variability in their entropies throughout the learning process. This variability could be due to the forgetfulness factor in CoACA, where all (*s, a*) pairs are multiplied by a *γ* (see Equation 4), reducing the value of the credits of all (*s, a*) pairs and thus increasing the uncertainty (and entropy) at the beginning of each session.

**Figure 4:**
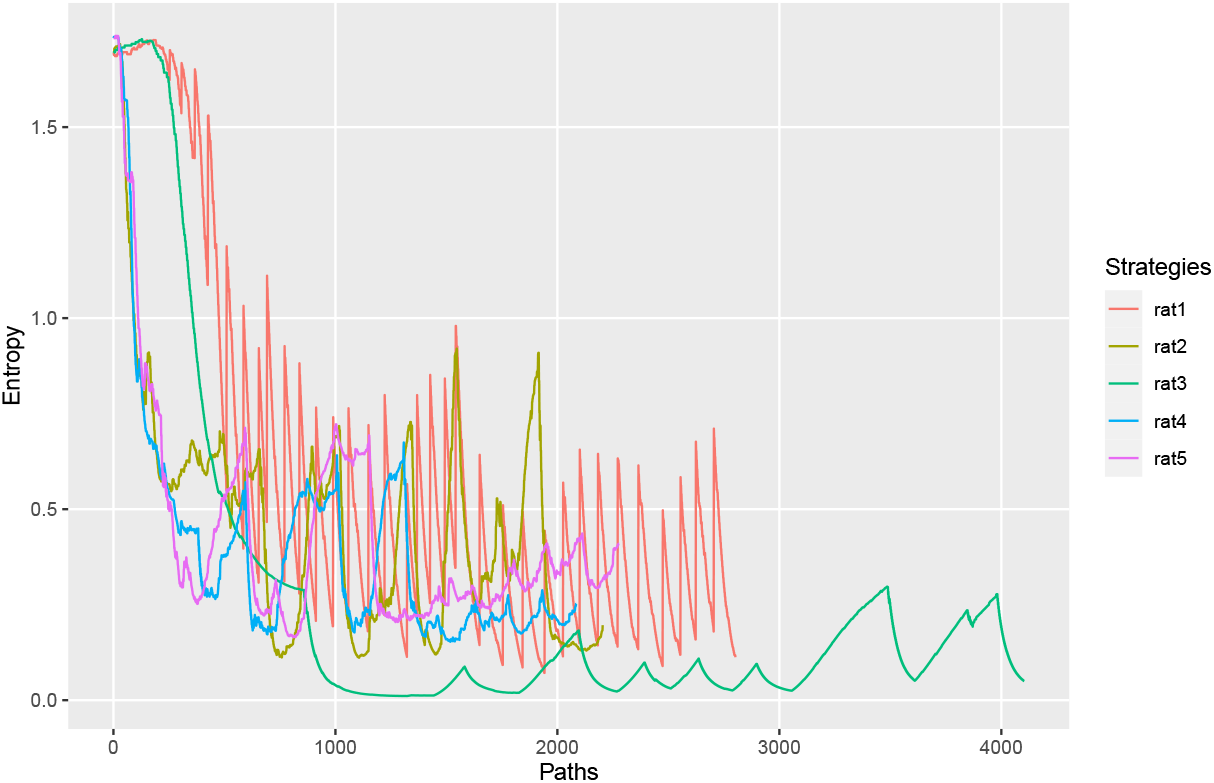
Entropy of rats’ policies for taking each path in the experiment, computed using the most likely agent model structure in Table 2 .

#### 2.5.1 Exploration *vs* Exploitation During Learning

In the context of the maze experimental task, the ideal behavior is for the rat to take the good path to claim a reward from one feeder and then take the good path again to go to the second feeder to claim the second reward, which would be the sequence: Good.LF,Good.RF,Good.LF, Good.RF…….

However, the rats do not immediately learn the two good paths from both feeders and exhibit suboptimal behavioral patterns during the initial sessions when they are still learning. Rats show varying degrees of uncertainty in their behavior during the learning process. Table 3 shows the longest continuous recurring pattern observed during the first 400 paths for each rat. Based on this, the behavior of the rats can be categorized into two distinct patterns:

**Table 3:**
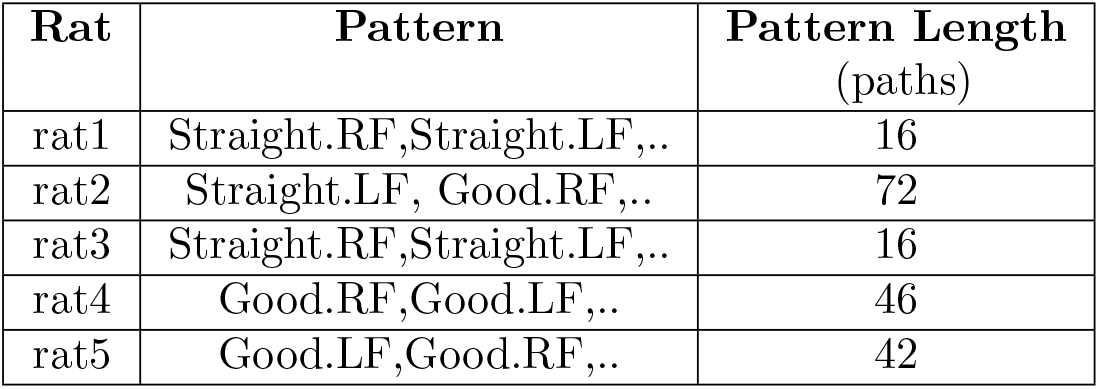
Longest continuous recurring pattern in the first 400 paths for each rat

| Rat | Pattern | Pattern Length<br>(paths) |
| --- | --- | --- |
| rat1 | Straight.RF,Straight.LF,.. | 16 |
| rat2 | Straight.LF, Good.RF,.. | 72 |
| rat3 | Straight.RF,Straight.LF,.. | 16 |
| rat4 | Good.RF,Good.LF,.. | 46 |
| rat5 | Good.LF,Good.RF,.. | 42 |

- Exploratory behavior: As shown in Table 3, rat1 and rat3 exhibit fewer consistent patterns during the initial sessions, with the longest sequence of Straight.LF and Straight.RF lasting only 16 paths in the first 400 paths. This indicates a high degree of inconsistency and uncertainty in their policy during the initial sessions inside the maze.
- Exploitative behavior: Rats 2, 4, and 5 exhibit more consistent behavior in the maze during the first 400 paths, indicating lower uncertainty in their policies. As shown in Table 3, rat2 learns only a partial reward path composed of Straight.LF and Good.RF during the initial stages but repeats it continuously for 76 paths, demonstrating exploitative behavior by attempting to repeat a partial reward solution. Rat4 and 5 learn both optimal paths early in the experiment and consistently repeat the ideal sequence to maximize rewards in the maze.

#### 2.5.2 Policy Entropy And Choice Durations

We assess whether there is any impact of policy entropy on the rats’ behavior during the course of the experiment. To examine the relationship between duration and entropy, a linear regression model was estimated using time series data. The model was specified as follows:

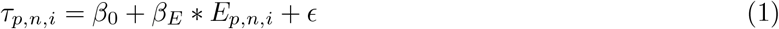

where duration is the dependent variable, entropy is the independent variable, *β*_0_ and *β*_*E*_ are the intercept and slope coefficients, respectively, and *ϵ* is the error term. *E*_*p*,*n*,*i*_ represents the policy entropy of *i*^*th*^ path of *n*^*th*^ episode of session *p*, computed using Equation 13 based on the most likely agent models from Table 2. *τ*_*p*,*n*,*i*_, the duration of the *i*^*th*^ path in episode *n* of session *p* is given by

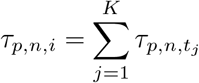

where *K* is the number of actions in *i*^*th*^ path of episode *n* of session *p* and 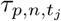 is duration of action 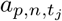 starting at time *t*_*j*_ in episode *n* of session *p*. To account for potential auto-correlation in the error terms, standard errors were adjusted using Newey-West method.

Table 4 presents the regression coefficients and their significance for each rat. The coefficient for entropy (*β*_*E*_) is positive for rats 1 and 5, indicating that these rats tend to increase their action duration as uncertainty (measured by entropy) rises. Conversely, rats 2 and 3 exhibit negative coefficients, suggesting they accelerate their actions under higher uncertainty. Rat 4 shows no significant relationship between entropy and action duration. Notably, these diverse responses to uncertainty do not appear to be linked to the rats’ internal agent model structures.

**Table 4:** Linear Regression Coefficients

| Rat | Agent Structure | $\beta_0$ | $\beta_E$ |
| --- | --- | --- | --- |
| rat1 | Hybrid3.CoACA | $\beta_0 = 4363.73, SE = 73.25,$<br>$pval < 2.2 \cdot 10^{-16}$ | $\beta_E = 340.69, SE = 103.24,$<br>$pval = 9.8 \cdot 10^{-4}$ |
| rat2 | Hybrid3.DRL | $\beta_0 = 3711.55, SE = 94.97,$<br>$pval < 2 \cdot 10^{-16}$ | $\beta_E = -410.78, SE = 170.46,$<br>$pval = 1.6 \cdot 10^{-2}$ |
| rat3 | Hybrid3.DRL | $\beta_0 = 3620.85, SE = 30.08,$<br>$pval < 2.2 \cdot 10^{-16}$ | $\beta_E = -199.52, SE = 61.67,$<br>$pval = 1.2 \cdot 10^{-3}$ |
| rat4 | Turns.DRL | $\beta_0 = 3463.89, SE = 125.02,$<br>$pval < 2 \cdot 10^{-16}$ | $\beta_E = 444.33, SE = 337.51,$<br>$pval = 0.1882$ |
| rat5 | Turns.DRL | $\beta_0 = 3283.73, SE = 62.05,$<br>$pval < 2.2 \cdot 10^{-16}$ | $\beta_E = 444.14, SE = 142.96,$<br>$pval = 1.9 \cdot 10^{-3}$ |

## 3 Discussion

This study aimed to illuminate the cognitive underpinnings of individual behavior, which can often deviate from rational reward-maximizing behavior, by investigating the interplay between learning rules and environmental representations. We defined that an RL agent’s internal model as a combination of both learning rule and environmental representation employed by individuals. Importantly, the optimality of the agent’s model is determined by the combination of the learning rule and environmental representation, rather than by each component individually. By systematically exploring a range of learning rules and environmental models, we sought to identify the optimal model structure for each individual. To identify the most probable internal model for each agent, we used the Bayesian Information Criterion (BIC) to assess the fit of various model configurations to the observed behavior.

We applied this structure learning framework to analyze rat behavior in a T-maze learning task where agent models were defined using two learning rules: the suboptimal CoACA, which favors actions with longer durations, and the optimal TD-learning-based Discounted RL (DRL), which maximizes expected returns. To model the rats’ internal environment representation, we defined six different cognitive graphs of the maze: Paths, Hybrid1, Hybrid2, Hybrid3, Hybrid4, and Turns. In total, we defined 12 distinct agent models for the maze task. We classified DRL with Turns model as the most efficient agent model structure in the maze task as it had more frequent TD updates compared to other agent models, which could speed up the learning in the task. CoACA-based agent models were classified as suboptimal due to their forgetfulness factor between sessions and the rest of the agent models were categorized as moderately optimal.

Experimental data revealed two distinct rat learning profiles: fast learners (rats 4 and 5) and slow learners (rats 1, 2, and 3) (see Figure 2b). Fast learners efficiently acquired optimal paths to both maze feeders, while slow learners demonstrated a more gradual learning process, often mastering one feeder’s path before discovering the other. The inferred agent models showed that slow-learning rats were best represented by suboptimal CoACA-based agent models and moderately optimal models, with CoACA and Hybrid3 being the best fit for rat 1, and DRL with Hybrid3 for rats 2 and 3. In contrast, DRL with Turns was the most suitable agent model structure for fast-learning rats. This confirms our hypothesis that Turns with DRL agent model due to its capacity for frequent updates, is essential for capturing the efficient learning behavior observed in the maze task.

Further qualitative analysis of individual behavior revealed additional intra-individual differences in the rats’ learning processes that were not fully captured by the inferred agent models. Rats 1 and 3 displayed more explorative behavior compared to the other rats during their learning phase. Additionally, the rats exhibited variable responses to high entropy (uncertainty): some rats, such as rats 1 and 5, increased their choice durations, while rats 2 and 4 decreased theirs.

Our approach of using structure learning to infer individual agent models—by examining their learning rules and internal environment representations—revealed intriguing aspects of rat behavior in the maze experiment. Although this study captures differences in individual behavior as a function of their learning rules and internal environment models, leading to suboptimal and optimal behaviors, it does not fully explain qualitative aspects such as variations in exploratory and exploitative behaviors among the rats. The current study assumes a stationary internal model for agents. Future research incorporating dynamic internal model switching could offer deeper insights into the qualitative aspects of individual decision-making.

Our research aimed to model the nuances of individual decision-making, including suboptimal behavior, by examining the interplay between an individual’s learning rule and their internal representation of their environment that define their internal cognitive agent models. Our findings suggest that suboptimal behavior is not solely a result of flawed learning rules or inaccurate environmental models (Reddy et al., 2018; Kanen et al., 2019; Lim et al., 2019; Zorowitz et al., 2020). Instead, the level of agent’s optimality depends on the specific combination of its internal model components, defined by its learning rule and internal environment representation, which may be optimal, suboptimal or moderately optimal in a given task. This insight offers a more sophisticated understanding of suboptimal behavior, framing it as an emergent property of an agent’s internal cognitive structure rather than a simple consequence of suboptimal learning process or a flawed perception of the environment. This perspective would also lead to the creation of more robust AI systems that have to interact with or recognize the intent of human subjects, even when they behave suboptimally. Such AI applications include explainable AI (xAI) (Arrieta et al., 2020; Beckers, 2022) and Human-Centered Artificial Intelligence (HCAI) (Riedl, 2019; Schmidt, 2020)), which are utilized in areas like pedestrian motion prediction (Ziebart et al., 2009), goal recognition (Meneguzzi and Pereira, 2021; Amado et al., 2022), human-robot interaction (Levine and Williams, 2014) among others.

## 4 Methods

### 4.1 Semi-Markov Decision Process

Temporal credit assignment can be represented as a Semi-Markov Decision Process (SMDP), which is a generalisation of a Markov Decision Process in which each action has a random duration. The SMDP can be thought of as a process in which individuals pass through *controlled* and *uncontrolled* states, where *controlled* states allow the individual to make decisions that influence the course of the process, whereas *uncontrolled* states do not allow the individual to make any decisions. The evolution of the system through uncontrolled and controlled states can be modelled as a *natural process* that keeps track of each state change in the system (Puterman, 2014; Parr, 1998).

An SMDP can be defined by a tuple

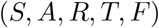

where *S* is the set of states, *A* is the set of actions, *R* is the reward function that gives the reward associated with each (*S, A*) in the environment, *T* is the transition function that gives the transition probabilities *P* (*s*′ | (*s, a*)), *F* : *F* (*t* |*s, a*), *t* ∈ ℝ^+^ gives the probability that next *controlled* state is reached within time *t* after action *a* is chosen in state *s*.

In the cognitive graphs of the maze models presented in Figure 3, the state *S* of the rat is defined by three variables: “last visited feeder box”, “last chosen action” and “current head position”. An action *A* is defined as choosing a trajectory at a given state and following that trajectory to reach the next state in the cognitive graph. The above state definition distinguishes actions based on both “last visited feeder box” and “last chosen action”. So, the action *A* → *B* → *RF* when coming from LF (*cf*. Figure 2a) is a different action to when coming from RF, and the action *B* → *RF* when coming from A is a different action to when coming from LF (cf. Figure 3).

### Episode In The Maze Task

An episode in the task is defined as a minimal segment of the rat’s trajectory where the rat starts from one feeder box, visits the other feeder box and returns to the starting box. Two examples of episode are given below, where 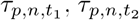 and 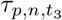 represents the durations of actions taken at times *t*_1_, *t*_2_ and *t*_3_ in episode *n* of session *p*:

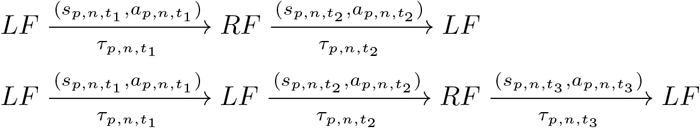

### Cognitive Activity-based Credit Assignment (CoACA)

CoACA is based on the idea of learning using *activity* from *Activity-based Credit Assignment*, where credit of action choices are accumulated through time. CoACA model is suitable for modeling situations where the assessment of choices incorporates both the environmental feedback (reward values) and the effort invested in making those choices. This distinguishes it from rational choice reinforcement learning models, which focus solely on future payoffs when making decisions. As a result, the treatment of duration differs between these models: reinforcement learning approaches view duration as a cost to be minimized to achieve maximum net benefits (returns), while in CoACA, duration reflects the effort invested in the choice. This is captured in CoACA using the idea of *activity*, which can be thought of as a way of measuring the cost/effort of an action.

In CoACA, *activity* is utilized to distribute the credit of rewards received during an episode among the actions selected within the episode, with greater *activity* resulting in greater credits. The rat learning problem, formulated as an SMDP where rats must recall state-defining variables such as “last visited feeder box” and “last chosen action”, provides an ideal setting for utilizing *activity* for credit allocation. In this context, *activity* can be seen as the expense of retaining the state variables in memory while executing an action. CoACA combines activity and reward to compute a credit that reflects effort (behavioral activity in an episode) of actions and the rewards obtained at the end of those episodes, which is accumulated over time. By virtue of the above properties, CoACA also implicitly rewards the repetition of successful past choices, even if they are suboptimal, and can model the behavior of rats when they persist with suboptimal action patterns.

Activity is computed as the duration of an action, relative to the duration of an episode:

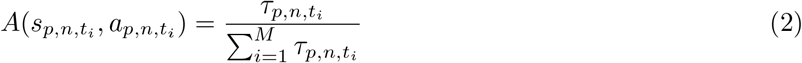

where *t*_*i*_ represents the the time of the *i*^*th*^ action in episode *n* of session *p*, where *i* ∈ [0, *M*] with M being the total number of actions in the *n*^*th*^ episode of *p*^*th*^ session. 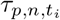 represents the duration of the action taken at time *t*_*i*_ in episode *n* of session *p*.

At the end of an episode *n* in session *p*, credits of all (*s, a*) selected during the episode are updated:

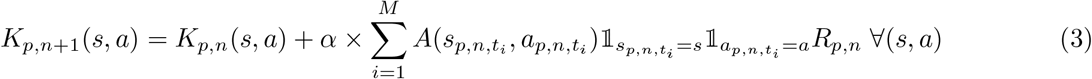

Here *t*_*i*_ represents the time at which *i*^*th*^ action of episode *n* in session *p* was taken, *i* ∈ [1, *M*], *R*_*p*,*n*_ = {0, 1, 2 } is the total reward obtained in episode *n* and *α* is the learning parameter (0, 1].

At the end of a session, the credits of all (*S, A*) pairs in the maze are decayed:

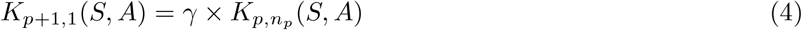

where *γ* ∈ [0, 1] is the forgetfulness parameter.

The probability of selecting an action *a* in state *s* is computed using the softmax rule:

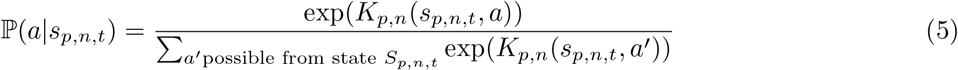

### Discounted Reward Q-learning

In discounted reward setting, a reward received in future is less valuable than an immediate reward. This means that the present value of one unit reward received *τ* ∈ R^+^ units in the future equals exp (*™ βτ*). The reward associated with transition from state *s*→ *s*′ is composed of two parts (Puterman, 2014)):

1. An immediate lump sum reward *L*(*s*_*t*_, *a*_*t*_).
2. Continuous reward accumulated at rate *ρ*(*W*_*t*_, *s, a*), where *W*_*t*_ represents the state of the *natural process* at time *t* :

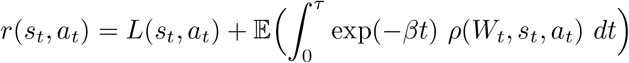 In the maze, the only reward is available at the end of an action duration. The value of future reward available after an action of duration *τ* is updated as:

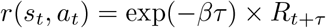

Where the lumpsum reward *L*(*s*_*t*_, *a*_*t*_) can be defined as decayed value of the reward *R*_*t*+*τ*_ obtained in the maze at the end of an action with duration *τ*, while the constant reward rate, *ρ* = 0.

SMDP Q-learning with discounted rewards uses temporal difference (TD) error to iteratively update Q-values (Bradtke and Duff, 1994).

Let 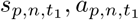 be part of episode *n* of session *p*, leading to new state 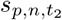 after duration 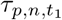 with a reward 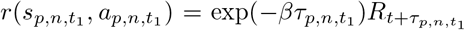 where 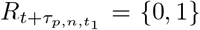 is the reward obtained in the maze after time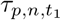, and *β* is the exponential discount factor applied to future rewards. This state transition can be noted as:

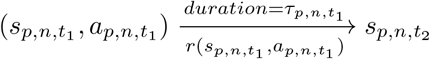

Since CoACA implicitly implements a memory trace of an episode, we implement an eligibility trace in DRL, lasting for the duration of a single episode:

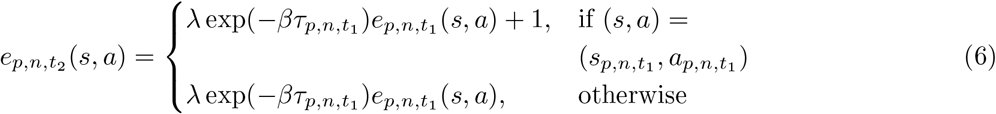

where 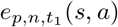 represents the eligibility trace of state-action pair (*s, a*) at time *t*_1_ in episode *n* of session *p*. At the end of an episode, *e*(*s, a*) = 0 ∀(*s, a*). Temporal difference prediction error *δ* is given by:

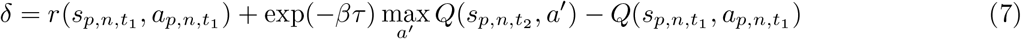

TD update is given by:

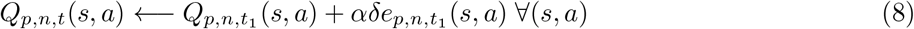

The probability of selection of action *a* at state *s*_*t*_ is

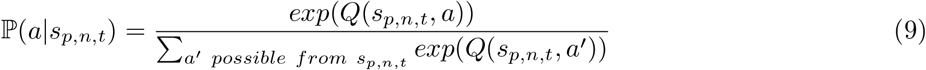

### 4.2 Model selection and validation

To compare how well the different models fit the data, the log likelihood of the model *m* is calculated using the model probabilities and is used as a measure of goodness of fit.

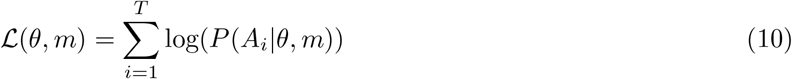

where *A*_*i*_ can be either *A*_*p*,*n*,*t*_ in case of DRL or CoACA, and *A*_*p*,*t*_ for ARL. Typically to avoid overfitting, likelihood measure of goodness-of-fit is penalized based on the number of model parameters (Daw et al., 2011; Wilson and Collins, 2019). Here we use Bayesian Information Criterion (BIC) (Schwarz, 1978) to penalize for the extra parameter in DRL based agent models, where BIC score of agent model *m* is computed as:

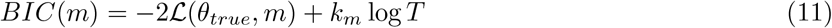

where *θ*_*true*_ is the agent model parameters estimated from rat’s experimental data by maximum likelihood estimation, *k*_*m*_ represents the number of parameters estimated in model *m* and *T* is the total number of paths between the two feeders in the rat’s experimental data. The parameter values of the models are determined from the experimental data using Maximum Likelihood Estimation (MLE) by maximizing the log likelihood of rats’ choices:

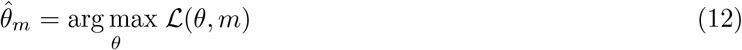

The parameter values that maximize the log likelihood (or minimizes the negative log likelihood) were determined using Self-adaptive Differential Evolution algorithm in Pagmo cpp package (Biscani and Izzo, 2020). The estimated model parameters are given in Supplementary Section 1.

## 5 Policy Entropy

In RL, policy *π*(*a*| *s*) is defined as the probability of taking action *a* in state *s*. Uncertainty in the policy corresponds to the randomness or variability in choosing actions given a particular state. The degree of uncertainty inherent in a policy reflects the variability in action selection for a given state. A deterministic policy, exhibiting low uncertainty, consistently chooses the same action in a particular state, indicating the agent has learned effectively. Conversely, a uniform policy, where all actions are equally likely, signifies high uncertainty in action selection, resulting in diverse behaviors within a state.

The entropy of the policy for the *i*^*th*^ path in episode *n* of session *p*, denoted as *E*_*p*,*n*,*i*_, is calculated as follows:

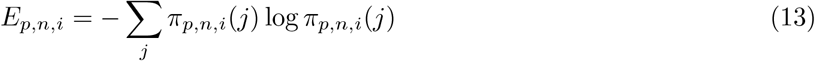

where *π*_*p*,*n*,*i*_(*j*) represents the probability of selecting path *j* as the *i*^*th*^ path of episode *n* in session *p*.

## Acknowledgements

This work was supported by the French government, through the UCA^Jedi^ and 3IA Côte d’Azur Investissements d’Avenir managed by the National Research Agency (ANR-15-IDEX-01 and ANR-19-P3IA-0002), by the interdisciplinary Institute for Modeling in Neuroscience and Cognition (NeuroMod) of the Université Côte d’Azur and by the National Research Agency (ANR-20-CE23-0004) with the DeepSee project. It is part of the Computabrain project.

NEF computing platform from Inria Sophia Antipolis Méditerranée Research Center (CRISAM) has been used for running or parallel simulations. NEF is part of the OPAL distributed computation mesocentre. The authors are grateful to the OPAL infrastructure from Université Côte d’Azur for providing resources and support.

## Footnotes

1 https://stat.ethz.ch/R-manual/R-devel/library/MASS/html/kde2d.html, last accessed on 08-19-2024

