## Supplementary Information for "Capturing Optimal and Suboptimal behavior Of Agents Via Structure Learning Of Their Internal Model"

September 30, 2024

### 1 Parameter settings

| Model | Parameter | Rat 1 | Rat 2 | Rat 3 | Rat 4 | Rat 5 |
|---|---|---|---|---|---|---|
| **Paths_aca2** | $\alpha$ | 0.0200 | 0.0282 | 0.0178 | 0.0186 | 0.0133 |
| | $\gamma$ | 0.4721 | 0.3476 | 0.4098 | 0.3245 | 0.4169 |
| **Hybrid1_aca2** | $\alpha$ | 0.0467 | 0.0428 | 0.0484 | 0.0392 | 0.0230 |
| | $\gamma$ | 0.3222 | 0.2829 | 0.2945 | 0.2014 | 0.3383 |
| **Hybrid2_aca2** | $\alpha$ | 0.0349 | 0.0426 | 0.0270 | 0.0326 | 0.0232 |
| | $\gamma$ | 0.4647 | 0.3701 | 0.4590 | 0.3145 | 0.4056 |
| **Hybrid3_aca2** | $\alpha$ | 0.0389 | 0.0452 | 0.0310 | 0.0365 | 0.0262 |
| | $\gamma$ | 0.4722 | 0.3767 | 0.4395 | 0.3144 | 0.4019 |
| **Hybrid4_aca2** | $\alpha$ | 0.0469 | 0.0409 | 0.0461 | 0.0380 | 0.0214 |
| | $\gamma$ | 0.2871 | 0.2621 | 0.2982 | 0.1831 | 0.3276 |
| **Turns_aca2** | $\alpha$ | 0.0701 | 0.0682 | 0.0536 | 0.0577 | 0.0369 |
| | $\gamma$ | 0.3657 | 0.3048 | 0.4196 | 0.2451 | 0.3644 |

Table 1: Model parameters for 5 different rats (Part 1). For ‘aca‘ models, parameters are: $\alpha$ (alpha) and $\gamma$ (gamma).

| Model | Parameter | Rat 1 | Rat 2 | Rat 3 | Rat 4 | Rat 5 |
|---|---|---|---|---|---|---|
| **Paths_drl** | $\alpha$ | 0.0052 | 0.0220 | 0.0110 | 0.0085 | 0.0071 |
| | $\beta$ | 0.0002 | 0.0002 | 0.0000 | 0.0003 | 0.0003 |
| | $\gamma$ | 0.9966 | 0.9990 | 0.8285 | 0.9999 | 0.9998 |
| **Hybrid1_drl** | $\alpha$ | 0.0182 | 0.0641 | 0.0118 | 0.0471 | 0.0465 |
| | $\beta$ | 0.0001 | 0.0001 | 0.0000 | 0.0001 | 0.0001 |
| | $\gamma$ | 0.9931 | 0.5352 | 0.9747 | 0.8724 | 0.8586 |
| **Hybrid2_drl** | $\alpha$ | 0.0273 | 0.0654 | 0.0145 | 0.0650 | 0.0658 |
| | $\beta$ | 0.0001 | 0.0001 | 0.0001 | 0.0001 | 0.0002 |
| | $\gamma$ | 0.5005 | 0.5932 | 0.8194 | 0.9045 | 0.6986 |
| **Hybrid3_drl** | $\alpha$ | 0.0222 | 0.0767 | 0.0163 | 0.0510 | 0.0529 |
| | $\beta$ | 0.0001 | 0.0001 | 0.0000 | 0.0001 | 0.0001 |
| | $\gamma$ | 0.7096 | 0.3553 | 0.7613 | 0.7790 | 0.8278 |
| **Hybrid4_drl** | $\alpha$ | 0.0238 | 0.0562 | 0.0135 | 0.0616 | 0.0576 |
| | $\beta$ | 0.0001 | 0.0001 | 0.0000 | 0.0001 | 0.0001 |
| | $\gamma$ | 0.9997 | 0.7777 | 0.9313 | 1.0000 | 0.9031 |
| **Turns_drl** | $\alpha$ | 0.0234 | 0.0801 | 0.0130 | 0.0574 | 0.0510 |
| | $\beta$ | 0.0001 | 0.0001 | 0.0001 | 0.0001 | 0.0002 |
| | $\gamma$ | 0.8906 | 0.6710 | 0.9209 | 0.9324 | 0.9694 |

Table 2: Model parameters for 5 different rats (Part 2). For ‘drl‘ models, parameters are: $\alpha$ (alpha), $\beta$ (beta), and $\gamma$ (gamma).
